# NeuID, a novel neuron-specific lncRNA, resolves a key epigenetic mechanism linking gene silencing to Alzheimer’s disease

**DOI:** 10.1101/2025.06.01.657217

**Authors:** Ranjit Pradhan, Zorica Petrovic, M. Sadman Sakib, Sophie Schröder, Dennis Manfred Krüger, Tonatiuh Pena, Eren Diniz, Susanne Burckhardt, Anna-Lena Schütz, Iga Grządzielewska, Karl Toischer, Thor D. Stein, Jan Krzysztof Blusztajn, Ivana Delalle, Jelena Radulovic, Farahnaz Sananbenesi, Andre Fischer

## Abstract

The increasing evidence that non-coding RNAs can become deregulated during pathogenesis is dramatically expanding the space for drug discovery beyond the protein-coding genome. Long noncoding RNAs (lncRNAs) are emerging as key regulators of cellular function, yet most remain uncharacterized. Here, we identify a previously unstudied lncRNA, which we named **Neu**ronal **Id**entity (*NeuID*)—a conserved, brain-enriched transcript expressed exclusively in neurons. *NeuID* is downregulated in the brains of Alzheimer’s disease (AD) patients. Mechanistically, *NeuID* maintains neuronal identity by repressing developmental and glial genes via interaction with the PRC2 subunit EZH2 and regulation of H3K27me3. Knockdown of *NeuID* disrupts this repression, leading to impaired neuronal activity and memory formation. Importantly, CRISPRa-mediated *NeuID* overexpression restores neuronal function in Aβ42-treated neurons. These findings identify *NeuID* as a critical regulator of neuronal plasticity and position it as a promising therapeutic target for AD.

**One sentence summary:** We identify *NeuID*, a novel brain and neuron-specific long non-coding RNA downregulated in Alzheimer’s disease, as a key regulator of neuronal identity and a promising therapeutic target to restore neuronal function.

## Introduction

Basic and translational research over the past decades has primarily focused on the protein-coding portion of the genome—that is, the ∼1.5% of genes translated into proteins and targeted by small molecules or biologics. In contrast, the vast majority of the genome is transcribed into non-coding RNAs (ncRNAs), including a substantial group known as long non-coding RNAs (lncRNAs), which are defined as transcripts longer than 300 nucleotides without protein-coding potential (*1, 2*). Initially dismissed as transcriptional noise, lncRNAs are now recognized as functional molecules involved in diverse biological processes (*3*) (*4*).

Many lncRNAs exhibit tissue- and cell type–specific expression patterns, with approximately 40% reported to be brain-specific (*5*). This suggests that lncRNAs play key roles not only in brain development and function, but potentially also in the pathogenesis of brain disorders (*6*) (*7*) (*8*) (*9*) (*10*) (*11*). These properties, alongside advances in RNA-based therapies, make lncRNAs attractive candidates for targeted drug development (*12*) (*13*).

Despite their promise, our understanding of lncRNAs in the brain—especially in the context of neurodegenerative diseases such as Alzheimer’s disease (AD)—remains limited (*14, 15*) (*16*).

AD is the most prevalent neurodegenerative disorder in the elderly and is believed to follow a specific sequence of pathological events that can be distinguished into different phases. While not undisputed, it is widely believed to begin with the accumulation of amyloid beta peptides, followed by aggregation of the Tau protein and deregulation of glial cells, eventually leading to loss of neuronal homeostasis and neuronal death (*17*) (*18*). Current therapeutic approaches focus on drugs that affect amyloid beta and Tau aggregation or modulate glial cell functions such as microglia-mediated neuroinflammation (*19*) (*20*) (*21*). However, AD is a multifactorial disease, and effective treatment will likely require combinatorial therapies using multiple approaches tailored to the patient’s disease stage (*22*).

On this basis, and in light of the emerging promise of RNA therapeutics for a wide range of diseases, we propose that a deeper understanding of lncRNAs in neuronal function can help develop precision therapeutic approaches to treat neurodegenerative diseases such as AD (*13*) (*23*) (*24*).

In this study, we systematically identified lncRNAs that fulfill these criteria. Using integrated OMICS datasets from humans and mice, we discovered a previously uncharacterized lncRNA, which we named ***Neu****ronal **Id**entity lncRNA* (NeuID). NeuID is almost exclusively expressed in neurons across species. Its expression is consistently reduced in the brains of AD patients and in cultured neurons exposed to Aβ42. Mechanistically, NeuID represses non-neuronal gene expression via the Polycomb Repressive Complex 2 (PRC2), which mediates transcriptional silencing through trimethylation of lysine 27 on histone H3 (H3K27me3). NeuID loss impairs neuronal plasticity and memory, while CRISPRa-mediated upregulation of NeuID restores network function in Aβ42-treated neurons.

In conclusion, our findings validate the feasibility of a systematic approach to identify lncRNAs as therapeutic targets in AD and highlight NeuID as a promising candidate for future RNA-based neuroprotective strategies.

## Results

### NeuID is a Conserved lncRNA Highly Enriched in Neurons of the Adult Brain

The primary objective of this study was to identify neuron-specific long non-coding RNAs (lncRNAs) that are enriched in brain tissue compared to other organs and to investigate their roles in Alzheimer’s disease (AD). We hypothesized that identifying such lncRNAs could reveal promising drug targets with minimal off-target effects, given their brain- and cell-type-specific expression. To test this, we integrated publicly available datasets with cell-type-specific total RNA sequencing (RNA-seq) and single-nucleus RNA sequencing (snRNA-seq) data from mouse and human brains **(Fig. 1a)**. The inclusion of both species was based on the premise that, particularly for lncRNAs, cross-species conservation is an indicator of functional relevance (*5*) (*25*).

**Figure 1.**
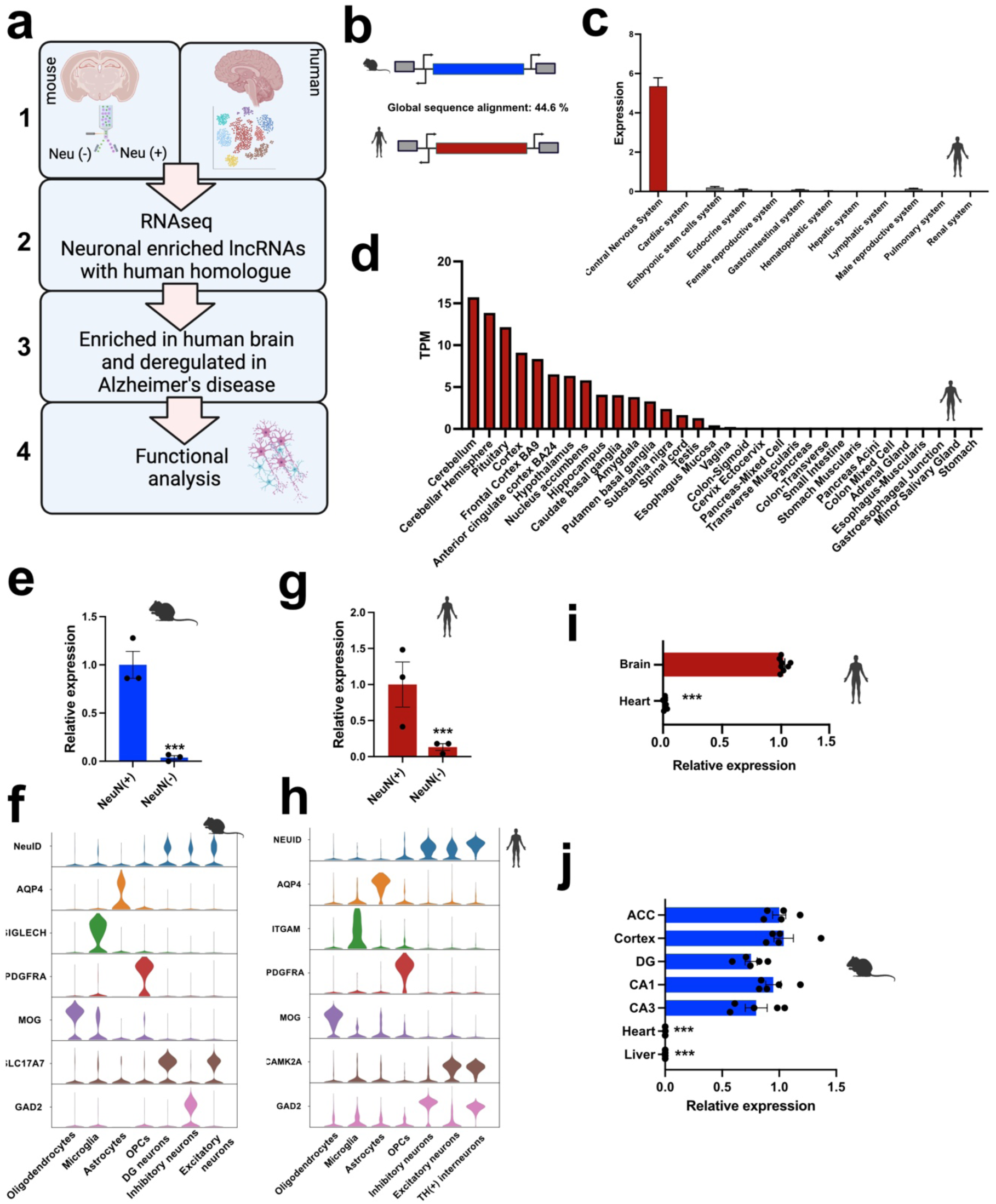
*NEUID* is a conserved lncRNA highly enriched in neurons of the adult brain. **a.** Schematic overview of the experimental approach. **b.** Schematic illustration showing the genomic localization of mouse *NeuID* and human *NEUID*. **c.** Expression of *NEUID* across various human tissues obtained from the RNA-atlas (*25, 27*), and **(d)** the GTEx tissue expression database (https://gtexportal.org). **e.** Bar chart showing qPCR analyses of *NeuID* expression between NeuN(+) (n=3) and NeuN (-) (n=3) nuclei obtained from the hippocampal CA1 region of mice. **f.** Violin plot depicting the expression of *NeuID* in a snucRNA-seq dataset from mice (*28*) brain along with marker genes for the depicted cell types. **g.** Bar chart showing qPCR analyses of *NEUID* expression in NeuN (+) (n=3) and NeuN (-) (n=3) nuclei from the human prefrontal cortex. **h)** Violin plot depicting the expression of *NEUID* in a snucRNA-seq dataset (*25*) obtained from the human prefrontal cortex along with marker gene expression or the depicted cell types. **i.** Bar chart showing qPCR analyses of *NEUID* in human brain (prefrontal cortex, n=8) and heart tissue (left ventricle, n=9). **j.** Bar chart showing qPCR analyses of *NEUID* expression in different mice brain areas (n=5), heart (n=3) and liver (n=4). ACC: Anterior cingulate cortex, DG: Dentate Gyrus, CA1/3: Cornu Ammonis 1/3. Asterisk indicates statistical significance analyzed via unpaired tTest. ***P < 0.0001. Error bars indicate SEM.

We began our analysis with mouse tissue by performing fluorescence-activated nuclear sorting (FANS) on the hippocampal CA1 region, a brain area essential for memory formation and among the first affected in AD. We isolated neuronal (NeuN⁺) and non-neuronal (NeuN⁻) nuclei from three-month-old wild-type mice and performed total RNA sequencing **(Fig. 1a, box 1)**. This analysis identified 247 lncRNAs that were significantly enriched in NeuN⁺ compared to NeuN⁻ nuclei (padj < 0.05, log₂FC > 3; **Supplemental Table 1**).

Next, we investigated whether any of these lncRNAs had human homologs **(Fig. 1a, box 2)**. We identified homologous lncRNAs based on shared syntenic genomic loci and a minimum sequence similarity of 40% (*26*) (*25*). Using this approach, we identified 15 candidate lncRNAs **(Supplemental Table 2)**. To further refine this selection, we leveraged a human RNA tissue atlas (*27*) (*25*) and the GTEx database (https://gtexportal.org) to filter for brain-enriched expression patterns **(Fig. 1a, box 3)**. This led us to a lncRNA, previously unstudied, which showed the highest enrichment in the human brain compared to other tissues **(Fig. 1b–d; Fig. S1)**. Given our later findings, we named this lncRNA ***Neu****ronal **Id**entity* (*NEUID*). In the following, *NeuID* refers to the mouse transcript, and *NEUID* to its human homologue.

To further validate the neuronal and brain-specific expression of *NeuID*, we performed qPCR analysis on NeuN⁺ and NeuN⁻ nuclei isolated from the mouse hippocampal CA1 region. Consistent with our sequencing data, qPCR confirmed that *NeuID* was exclusively expressed in NeuN⁺ nuclei **(Fig. 1e)**. Additionally, single-nucleus RNA sequencing (snRNA-seq) of mouse brain tissue confirmed that *NeuID* is highly enriched in neurons compared to glial cells **(Fig. 1f)**.

To determine whether a similar expression pattern exists in the human brain, we performed FANS on NeuN⁺ and NeuN⁻ nuclei isolated from the prefrontal cortex of individuals without neurodegenerative disorders. qPCR analysis revealed that *NEUID* was highly enriched in NeuN⁺ nuclei, mirroring our findings in mice **(Fig. 1g)**. Furthermore, snRNA-seq analysis of the human prefrontal cortex confirmed that *NEUID* is significantly enriched in neurons compared to glial cells **(Fig. 1h)**. Please note that the differences in expression levels detected via snucRNA-seq in mouse and human samples should not be directly compared, as the mouse data were generated using the 10X Genomics platform (*28*), while the human data were obtained using a modified protocol based on the Takara iCELL8 system, which was specifically adapted to enhance the detection of lncRNAs (*25*).

Finally, we further validated these findings in bulk tissue by performing qPCR analysis on total RNA isolated from the human prefrontal cortex and heart (left ventricle). The heart was chosen since, next to the brain, it is the only other organ that mainly consists of postmitotic and excitable cells. *NEUID* expression was significantly higher in the brain, with negligible expression detected in the heart **(Fig. 1i)**.

We also assessed *NeuID* expression across different mouse brain regions using qPCR and compared it to expression levels in the heart and liver. *NeuID* was consistently highly expressed in all analyzed brain regions, with minimal or undetectable expression in the heart and liver **(Fig. 1j)**.

In summary, we identified *NEUID* as a previously uncharacterized lncRNA with a neuron-specific and brain-enriched expression pattern in both mice and humans. Next we wanted to test if *NEUID* is deregulated in neurodegenerative diseases **(Fig. 1a**, **Box 4).**

### NEUID is down-regulated in the brains of AD patients

Given the emerging role of ncRNAs as novel drug targets in neurodegenerative diseases, we investigated whether *NEUID* is dysregulated in the brains of AD patients. Analysis of data from the AGORA database, which includes over 1,000 brain samples from healthy individuals and AD patients, revealed that *NEUID* expression is significantly reduced in the parahippocampal gyrus (PHG), dorsolateral prefrontal cortex (DLPFC), and temporal cortex (TCX) of AD patients compared to controls **(Fig. 2a)**.

**Figure 2.**
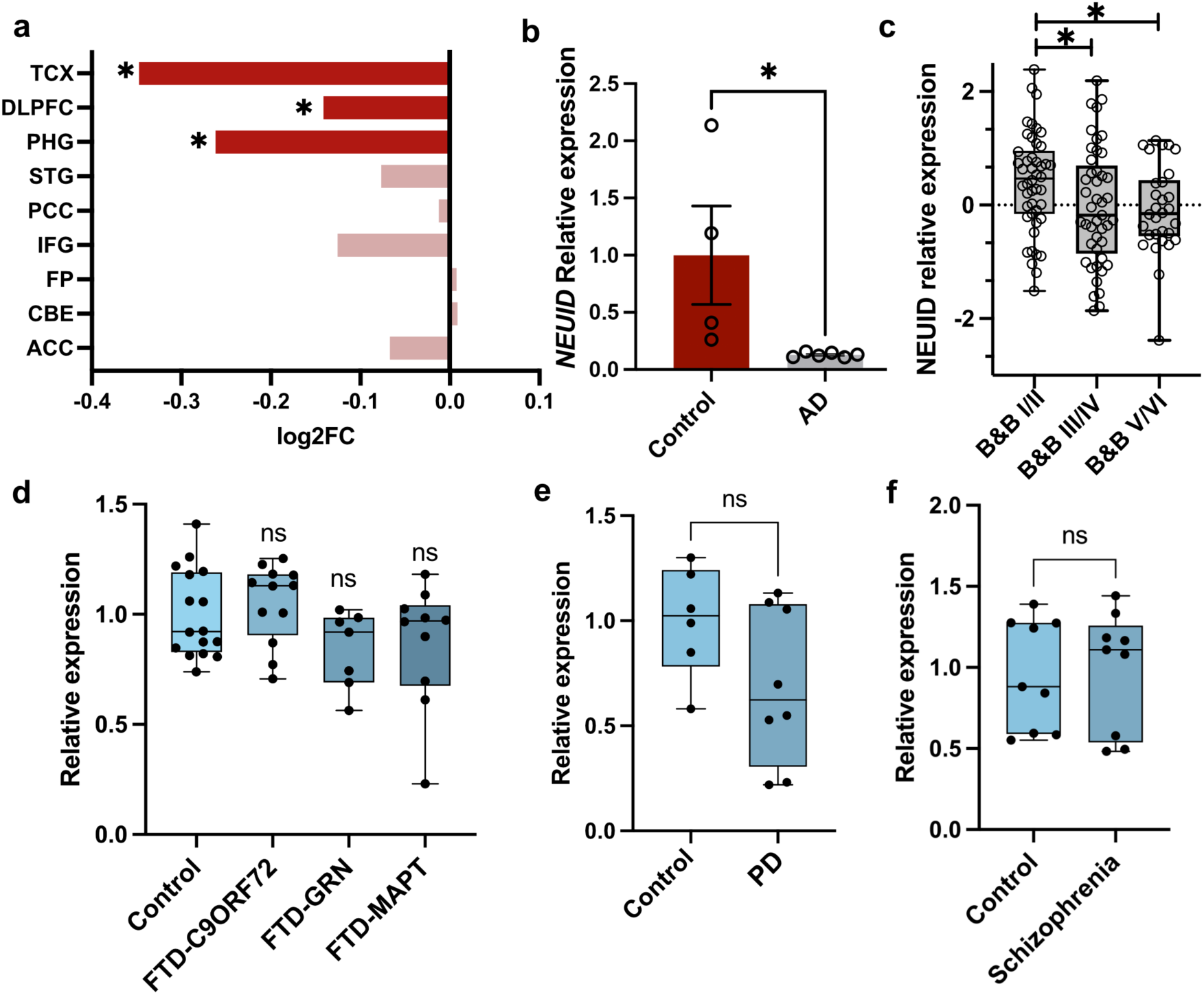
*NEUID* is down-regulated in the brains of AD patients. **a.** *NEUID* expression in various brain regions of AD and control postmortem human brains. Data were obtained from the AGORA AD database (\**p* < 0.05). **b.** qPCR analysis of *NEUID* expression in prefrontal cortex (Brodmann Area 9, BA9) postmortem brain tissue from control (n = 4) and AD (n = 6) patients (Unpaired tTest ; \**p* < 0.05). **c.** Expression of *NEUID* in postmortem brain tissue from individuals spanning Braak & Braak (B&B) stages I–VI. Data were obtained from the Framingham Heart Study (One way ANOVA; *P* = 0,034). **d.** Box plot showing *NEUID* expression in the frontal cortex of FTLD patients with mutations in C9ORF72 (n = 12), GRN (n = 7), MAPT (n = 10), and controls (n = 16) (*ns = not significant*). **e.** Box plot of *NEUID* expression in the frontal cortex of Parkinson’s disease (PD) patients (n = 8) and controls (n = 6). Data were obtained from the GEO dataset GSE156928 (*ns = not significant*). **f.** qPCR analysis of NEUID expression in the prefrontal cortex (BA9) of schizophrenia patients (n = 9) and controls (n = 9) (*ns = not significant*). Abbreviations: TCX, temporal cortex; DLPFC, dorsolateral prefrontal cortex; PHG, parahippocampal gyrus; STG, superior temporal gyrus; PCC, posterior cingulate cortex; IFG, inferior frontal gyrus; FP, frontal pole; CBE, cerebellum; ACC, anterior cingulate cortex; FTD, frontotemporal dementia; PD, Parkinson’s disease.

To confirm this observation, we performed qPCR analysis on a smaller cohort of postmortem brain samples characterized by Braak & Braak (B&B) stage IV. B&B staging is a widely used classification system for assessing the severity of AD pathology. B&B stages range from stage 1, representing minimal neurofibrillary tangle deposition, to stage 6, which indicates advanced, widespread pathology (*29*). In agreement with the AGORA findings, *NEUID* expression was significantly decreased in the prefrontal cortex (B&B stage IV, representing an early to moderate disease stage in the prefrontal cortex) when compared to controls **(Fig. 2b)**.

We further analyzed *NEUID* expression using the previously-described RNASeq dataset from brain tissue (prefrontal cortex, area 9) from the Framingham Heart Study (*30*) (*31*), focusing on samples spanning all B&B stages. Consistent with our prior results, *NEUID* expression was negatively correlated with B&B stages (r = 0,31; *P* = 0,049), indicating a progressive decline as pathology worsens **(Fig. 2c)**.

Interestingly, no significant changes in *NEUID* expression were detected in RNA-seq datasets derived from postmortem brain tissue of patients with frontotemporal lobar degeneration (FTLD) (*32*) (*33*) **(Fig. 2d)**, Parkinson’s disease (PD) (GEO dataset GSE156928) **(Fig. 2e)**, or schizophrenia (*34*) **(Fig. 2f)**. This indicates that *NEUID* dysregulation may be uniquely associated with AD pathology and moreover suggest that decreased *NEUID* levels are not simply a result of neuronal cell death.

### Knock down of NeuID affects neuronal gene-expression

To the best of our knowledge, *NeuID* has not been studied in any cellular context. Given our data showing that *NEUID* is enriched in neurons, almost exclusively expressed in the brain, and decreased in the brains of AD patients, we sought to elucidate its function.

Since lncRNAs perform regulatory functions specific to their cellular localization (*35*), we first assessed the localization of *NeuID*. To this end, we performed RNAscope in adult mouse brain sections, combined with NEUN staining to label neuronal nuclei. Our data confirmed that *NeuID* is highly expressed in neurons and further suggested that *NeuID* is enriched in neuronal nuclei **(Fig. 3a)**. Interestingly, two to ten bright foci of *NeuID* of different size were observed per neuronal nucleus, a pattern consistent with previous lncRNA studies and has been associated with the regulation of chromatin dynamics (*36*) (*37*). Consistent with this observation, qPCR analysis revealed that *NeuID* is enriched in nuclear fractions isolated from the mouse hippocampus compared to cytoplasmic fractions **(Fig. 3b)**, further suggesting a potential role in gene expression regulation.

**Figure 3.**
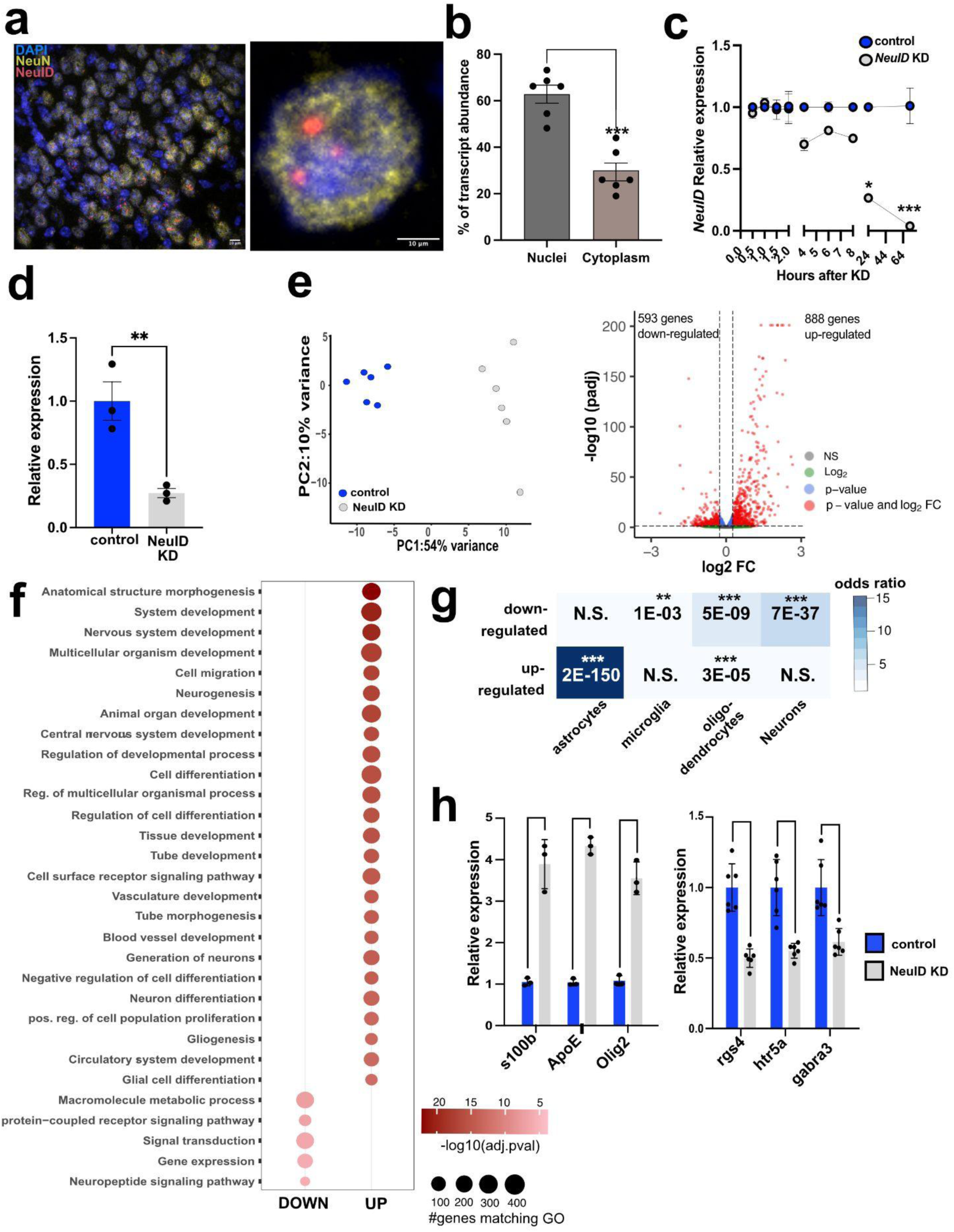
Knock down of NeuID affects neuronal gene-expression. **a.** Representative images from the adult mouse hippocampus showing RNAscope signal for NeuID (red) and neuronal nuclei stained with NeuN (green). DAPI was used to stain nuclei (blue). Scale bar: 10 µm. **b.** RT-qPCR quantification of NeuID expression in nuclear and cytoplasmic fractions of hippocampal neurons (n=6). unpaired tTest; ***p < 0.001. **c.** RT-qPCR quantification of NeuID expression at indicated time points following administration of NeuID Gapmers or control Gapmers in hippocampal neurons. One-way ANOVA revealed a significant group difference (*P* < 0.0001, F = 16.91); *p < 0.05, ****p < 0.001. **d.** Bar plot showing NeuID knockdown in RNA-seq samples (n=3). **p < 0.01, unpaired tTest. **e. Left panel:** Principal component analysis (PCA) of the RNAseq data**. Right panel:** Volcano plot depicting differentially expressed genes following NeuID KD (*FDR* < 0.05, |log2FC| > 0.5). **f.** Dot plot showing the top 10 % Gene Ontology (GO) terms (biological processes) associated with upregulated and downregulated genes after NeuID KD. **g.** Heatmap showing cell-type specific enrichment of the up- and downregulated genes. *p < 0.05, **p < 0.001, ***p < 0.0001, (hypergeometric test). **h.** Bar chart showing RT-qPCR quantification of selected genes de-regulated upon *NeuID* KD in the RNA-seq data(n=3 & n=6). Unpaired tTest.

To investigate *NeuID’s* function, we performed *NeuID* knockdown (KD) in primary hippocampal neurons using Gapmers, single-stranded antisense oligonucleotides that mediate RNase H-dependent degradation of nuclear RNA transcripts. As a negative control, we used Gapmers with no known target in the genome.

First, we assessed the efficacy of *NeuID* knockdown by qPCR analysis at multiple time points (0.5, 1, 1.5, 2, 4, 6, 7, 25, and 72 hours) after Gapmer administration. *NeuID* expression was fully suppressed 72 hours post-treatment **(Fig. 3c)**, and we used this time point for RNA sequencing (RNA-seq) to determine the effects of *NeuID* KD on the hippocampal neuronal transcriptome. *NeuID* knockdown efficacy was confirmed by qPCR **(Fig. 3d)**. RNA-seq analysis revealed 888 upregulated and 593 downregulated genes following *NeuID* KD (false discovery rate [FDR]; padj < 0.05, FC > 1.2 or < -1.2; **Fig. 3e**). Gene Ontology (GO) term analysis showed that upregulated genes were enriched in biological processes (**supplemental table 3**) related to development and glia cell function including the GO term such as “regulation of developmental processes”, “nervous system development”, “gliogenesis” and “glia cell differentiation **(Fig. 3f)**. For the down-regulated genes we detected much less significant GO-terms, when compared to the up-regulated genes including the GO terms “signal transduction”, “gene expression” and “neuropeptide signaling pathway” **(Fig 3f, supplemental table 4).** These data suggest that loss of *NeuID* leads to an orchestrated increased expression of glial cell genes and genes linked to developmental processes. In line with this interpretation we observed that the upregulated genes were enriched for genes normally expressed in astrocytes and oligodendrocytes, while the down-regulated genes were obviously highly enriched for neuronal genes **(Fig. 3g)**.

To validate these RNA-seq findings, we confirmed the altered expression of selected genes using qPCR. Our data show increased expression of non-neuronal genes, including oligodendrocyte transcription factor 2 (*Olig2*), which is normally expressed in oligodendrocytes; S100 calcium-binding protein B *(S100B*) normally expressed in astrocytes and apolipoprotein E (*ApoE*), normally expressed in astrocytes and microglia **(Fig. 3g)**. Conversely, genes such as gamma-aminobutyric acid type A receptor subunit alpha-3 (*Gabra3*), regulator of G-protein signaling 4 (*Rgs4*), and 5-hydroxytryptamine receptor 5A (*Htr5a*) - whose loss of function has been linked to cognitive impairments (*38*) (*39*) (*40*) - were downregulated **(Fig. 3g)**. These data indicate that *NeuID* plays a role in the orchestration of neuron-specific gene expression, as its reduced levels lead to increased expression of non-neuronal genes linked to developmental glial-related processes. This is paralleled by the downregulation of neuronal genes linked to memory function.

### NeuID regulates neuronal activity, synapse number and memory formation in mice

The analysis of neuronal gene-expression upon *NeuID* KD suggests that *NeuID* may play an important role in the regulation of neuronal functions and the loss of *NeuID* expression, as observed in AD patients, may cause deregulation of neuronal plasticity. To address this directly we decided to study neuronal network plasticity upon *NeuID* KD. Therefore, primary hippocampal neurons were subjected to electrical recordings using multi-electrode arrays (MEA). Hippocampal neurons were plated on MEA plates and *NeuID* KD was performed on DIV7. The MEA activity was recorded 72h later at DIV10 for 15 minutes.

Loss of *NeuID* resulted in severe impairments of neuronal network activity as indicated by a reduced neural activity score (NAS), a composite score that serves as an objective metric to compare neuronal activity in MEA (*41*) **(Fig 4a)** and a significantly impaired mean firing rate **(Fig 4b)**. These data suggest that neuronal network function is impaired when *NeuID* levels are reduced. Therefore, we further examined the role of *NeuID* in synchronized network burst activity. *NeuID* KD led to sparse electrical activity **(Fig 4c)** and decreased frequency of network bursts **(Fig 4d)**. In line with these data, the synchronous firing of neurons was also decreased in *NeuID* KD neurons when compared to the control group **(Fig 4e)**. These results indicate that decreased levels of *NeuID* impaired spontaneous neuronal firing and impedes synchronized network activity. One possibility to explain these data could be that *NeuID* might be important for maintaining the number and function of physiological synapses within a neuronal network. To test this hypothesis, we employed immunohistochemical approaches to measure synaptic markers informing about the density and number of synapses in response to *NeuID* KD. Our analyses revealed that KD of *NeuID* decreased dendritic spine density **(Fig 4f)** as measure by DIL dye staining, and led to a reduction in synapse numbers as measured by synaptophysin (Syn) and postsynaptic density 95 (PSD95) costainging **(Fig 4g)**.

**Figure 4.**
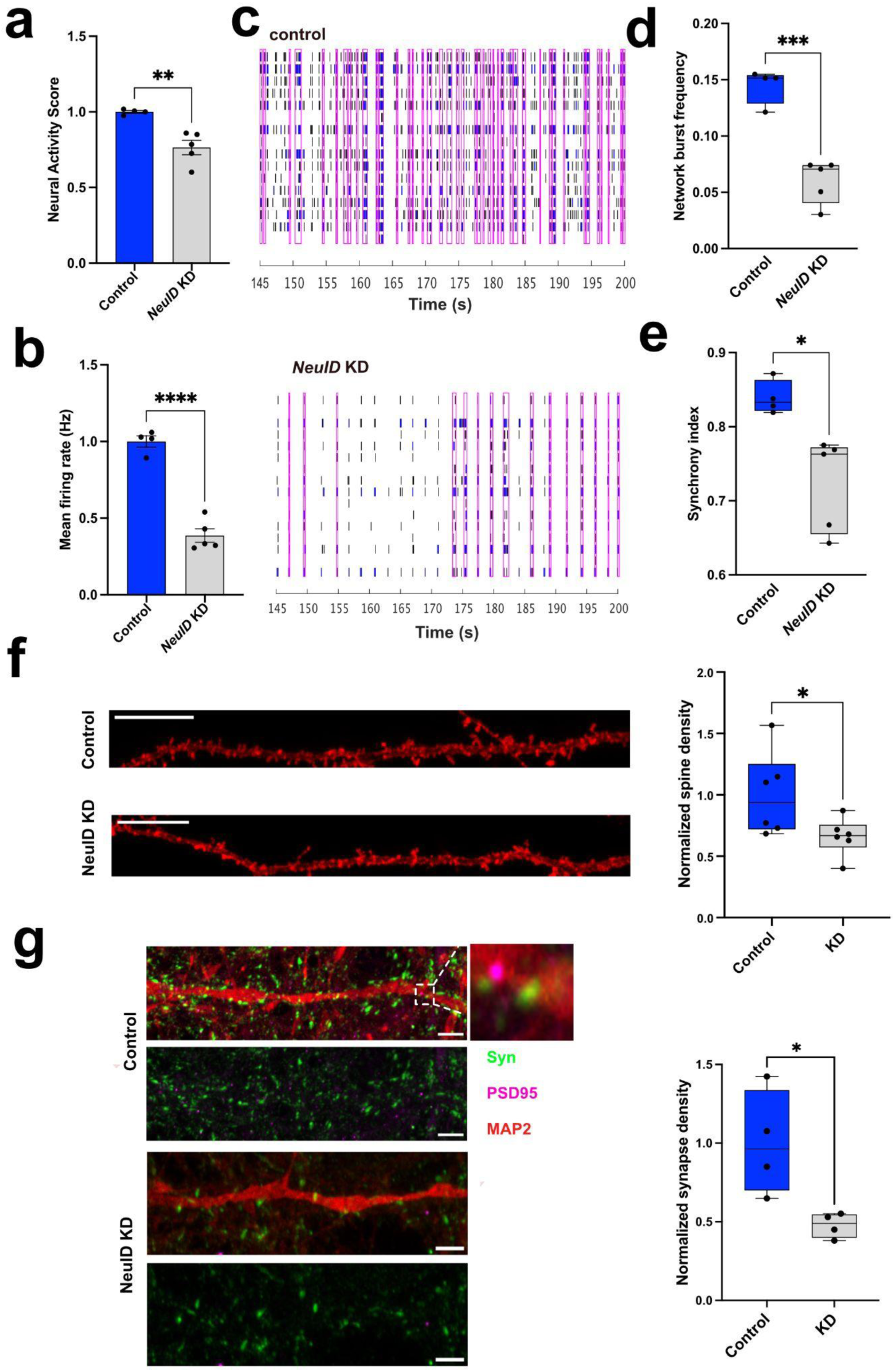
NeuID regulates neuronal activity through modulation of synapses. **a.** Bar chart displaying the NAS measured via an MEA assay. *p < 0.05 NeuID KD (n=4) vs control (n=5) ; Unpraired tTest. **b.** Bar plot showing the mean firing rate in control (n=4) and NeuID KD (n=5) conditions. ****p< 0.0001;Unpaired tTest. **c.** Representative raster plots showing neuronal network activity in control (n=4) and NeuID KD (n=5) neurons. **d.** Box plot quantifying network burst frequency amongst groups. ***p<0.001 NeuID KD vs control; Unpaired tTest. **e.** Box plot showing the synchrony index of control vs. NeuID neurons. *p < 0.05 NeuID KD vs control; Unpaired tTest. **f. Left panel:** Representative images showing traced neuronal dendrites stained with DiI dye in control and NeuID KD conditions **Right panel:** Box plot showing normalized dendritic spine density. *p < 0.05 NeuID KD vs control; Unpaired tTest. **g. Left panel:** Representative images of synapse quantification via IHC using synaptic markers synaptophysin (syn) and postsynaptic density (PSD-95). Neuron processes are stained with Map2. **Right panel:** Box plot showing the normalized synapse density. *p < 0.05; NeuID KD vs control; Unpaired tTest. Hz, Herz.

Collectively, these data support the view that *NeuID* plays a key role in the regulation of neuronal activity.

### Decreased levels of NeuID impair memory formation in mice

Our findings suggest that *NeuID* might affect cognitive processes such as memory consolidation. To test this possibility directly, we employed stereotactic injections to administer *NeuID* Gapmer’s into the dorsal hippocampus of wild-type mice and subjected them to a battery of behavioral tests to assess hippocampus-dependent memory formation. To this end, a group of mice was injected into the hippocampus with *NeuID* Gapmer’s or control oligomers, and hippocampal tissue was isolated eight days later **(Fig. 5a)**.

**Figure 5.**
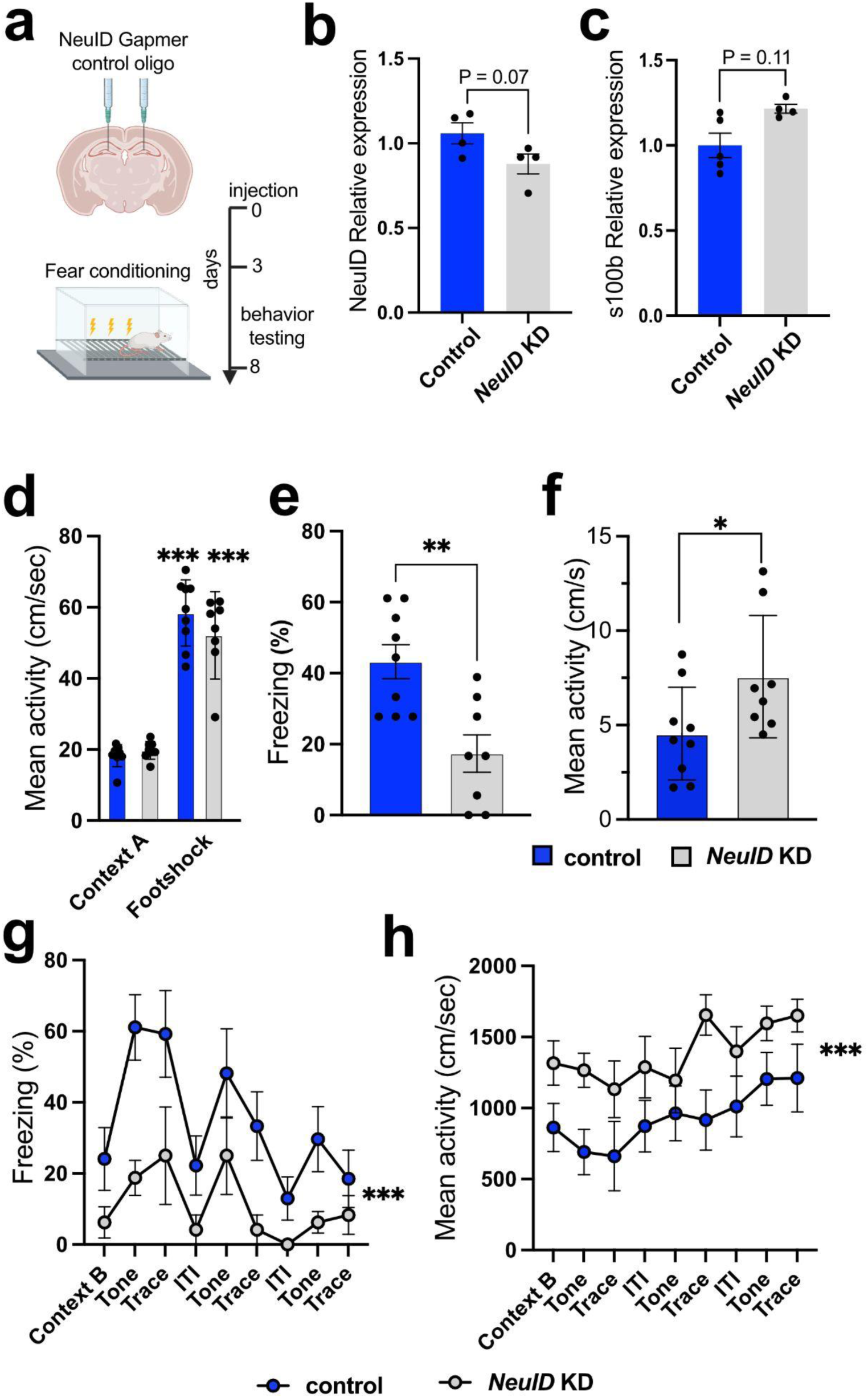
Decreased levels of *NeuID* impair memory formation in mice. **a.** Schematic depiction of experimental design. **b.** Bar chart showing qPCR data for *NeuID* expression in the hippocampus of mice, 8 days after the injection of scrambled oligomers (control, n=4) or *NeuID* Gapmer’s (*NeuID* knock down (KD, n=4), P = 0.0788, unpaired tTest. **c.** Bar chart depicting the expression of *s100b* in the hippocampus of mice treated as described for (b). P = 0.11, unpaired tTest. **d.** Bar chart depicting the mean activity during the fear conditioning training, indicating that both groups responded similarly to the foot shock (Control, n=9; NeuID KD, n=8). ***P < 0.0001 for Context A vs Footshook, unpaired tTest. **e.** Bar chart depicting freezing behavior during the memory test (re-exposure to context A). **P < 0.01, unpaired tTest. **f.** Bar chart depicting the mean activity during the memory test (re-exposure to context A). *P < 0.05, unpaired tTest. **g.** Freezing behavior and mean activity **(h)** during the memory test for trace fear conditioning. Context B: freezing during re-exposure to context B without presentation of the tone; Tone: freezing during re-exposure to context B during the presentation of the tone; ITI: freezing during the ITI (inter trial interval). ***P < 0.0001, Two-way ANOVA for control vs *NeuID* KD. Error bars indicate SD.

qPCR analysis revealed decreased *NeuID* levels in mice that received *NeuID* Gapmers compared to the control group, although statistical significance was only reached when applying FDR < 0.1 **(Fig. 5b)**. We also analyzed the expression of *S100b*, a gene that was upregulated following NeuID knockdown in primary hippocampal neurons (see Fig. 3h). Consistent with these findings, we observed a trend toward increased *S100b* levels in the hippocampus of mice treated with Gapmers; however, this change did not reach statistical significance (P = 0.11; **Fig. 5c**).

It is important to note that the reduction in *NeuID* and increased expression of *s100b* was less pronounced compared to the data obtained from neuronal cultures. This is likely due to the fact that not all hippocampal neurons are affected by stereotactic injection. However, previous studies have demonstrated that hippocampus-dependent memory impairment is detectable even when genes essential for memory formation are disrupted in as little as 10–20% of all hippocampal neurons (*42*).

Therefore we injected another group of mice into the dorsal hippocampus with either *NeuID* Gapmers or control oligonucleotides. Three days later, we subjected them to a fear conditioning paradigm which allows the measurement of contextual and trace fear conditioning, both of which are well-established hippocampus-dependent associative learning tasks **(Fig. 5a)**.

During the training session, activity levels and responses to the electric foot shock were similar between groups **(Fig. 5d)**. Freezing behavior, a widely used measure of memory consolidation, was significantly impaired in *NeuID* Gapmer-treated mice when tested 24 hours after training. The memory test consisted of a re-exposure to the conditioning context (Context A), during which control mice displayed robust freezing, whereas mice injected with NeuID Gapmers showed significantly reduced freezing behavior **(Fig. 5e)**. This impairment was accompanied by increased locomotor activity during the test session, which is in agreement with the reduced freezing behavior **(Fig. 5f)**.

We observed similar results in the trace fear conditioning test. During training, mice were placed in Context A, and after three minutes, exposed to a tone for 30 seconds followed by a 15-s trace period during which no stimuli were presented, terminating with a 2-s electric footshock. The tone-trace shock sequence was repeated 3 times, separated by 60-s intertrial (ITI) intervals. Memory recall was tested 24 hours later by placing the mice into a novel environment (Context B) and presenting the tone. During the test, freezing behavior significantly increased in Context B when the tone was played and during the trace period, but decreased during the inter-trial interval (ITI), when no stimulus was present.

*NeuID* Gapmer-treated mice exhibited a significantly impaired freezing response throughout the memory tests **(Fig. 5g)**, indicating deficient associative memory. This impairment was again accompanied by increased locomotor activity **(Fig. 5h)**, further supporting the hypothesis that *NeuID* KD disrupts hippocampus-dependent memory formation.

### CRISPR-mediated overexpression of NeuID rescues Aβ-mediated loss of neuronal plasticity

In summary, our findings provide strong evidence that physiological expression levels of *NeuID* are essential for neuronal plasticity and memory function, while decreased levels are observed in patients who develop AD. Based on this, we hypothesized that *NeuID* could be a novel drug target for AD.

To test this, we employed neurons treated with amyloid beta (Aβ) 42 oligomers as a model system, since our data suggest that *NeuID* expression decreases early in disease progression, and amyloid pathology is widely accepted as one of the earliest molecular changes observed in AD patients (*43*). Therefore, we first investigated the effect of Aβ42 exposure on *NeuID* expression in primary hippocampal neurons. qPCR analysis, performed 24 hours after treatment, revealed that *NeuID* levels were significantly decreased in Aβ42-treated neurons compared to the vehicle control group **(Fig 6a)**. To confirm the efficacy of Aβ42 treatment and establish a relevant read out for amyloid pathology, we assessed neuronal activity via MEA recordings. We first confirmed that neuronal activity was similar across all cultures before hippocampal neurons were treated with Aβ42 or vehicle control at DIV10. Recordings were performed 24 hours later for 15 minutes. The data show that Aβ42 treatment led to sparse electrical activity **(Fig 6b)**. In agreement with this, the NAS **(Fig 6c)**, the mean firing rate **(Fig 6d)** and the frequency of network bursts **(Fig 6e)** were significantly reduced upon Aβ42 treatment. These results show that exposure to Aβ42 leads to reduced *NeuID* levels and decreased neuronal activity.

**Figure 6.**
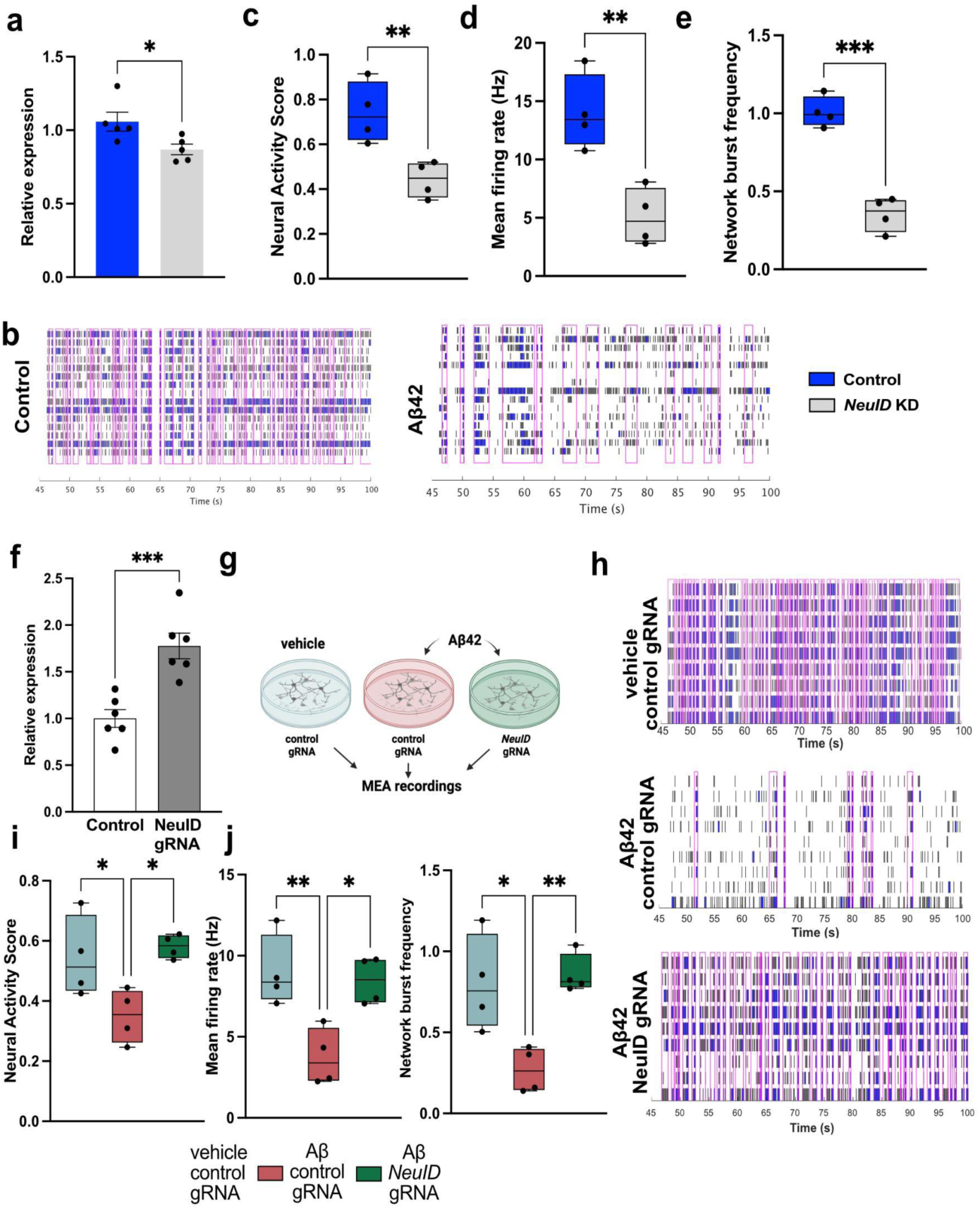
CRIPRa mediated over-expression of *NeuID* rescues Aβ mediated neuronal impairment. **a.** Bar plot showing the relative expression of *NeuID* in primary hippocampal neurons after treatment with Aβ42 (n=5); unpaired tTest *p< 0.05. **b.** Representative raster plots showing electrical activity during MEA recordings in the control and Aβ42 treated groups (n=4). **c.** Box plots showing the NAS, **(d)** the mean firing rate and **(e)** the network burst frequency of hippocampal neurons without (control) and after Aβ42 treatment. **f.** Bar plots showing expression of *NeuID* in hippocampal neurons treated with CRISPRa and NeuID gRNA (NeuID gRNA, n=6) and cells that were treated with a control gRNA (control, n=6). **g.** Experimental scheme for the experiments shown in h-j. **h.** Representative raster plots showing electrical activity during MEA recordings of hippocampal neurons that were treated with CRISPRa and either a control gRNA + vehicle (vehicle control gRNA), Aβ42 + control gRNA (Aβ42 control gRNA) or Aβ42 along with the *NeuID* gRNA (Aβ42 NeuID gRNA,). **i.** Box plots showing the NAS, **(j)** the mean firing rate and **(e)** the network burst frequency of hippocampal neurons subjected to the experiment described in (g) (n=4).*p<0.05, **p< 0.01, ***p< 0.001; unpaired tTest.

Having established that MEA recordings provide a suitable readout for amyloid pathology in our experimental setting, we next sought to determine whether increasing *NeuID* levels could reverse these phenotypes. To achieve *NeuID* overexpression, we used the CRISPR-based activation (CRISPRa) method. We generated a guide (g) RNA targeting the *NeuID* promoter, along with a corresponding control gRNA with no target in the genome (*25*). Upon transfection of the CRISPRa plasmid and gRNA into primary hippocampal neurons, we observed a twofold increase in *NeuID* expression **(Fig 6f)**.

Next, we transfected neurons with CRISPRa carrying either the *NeuID* or control gRNA and subjected them to Aβ42 treatment. Neurons transfected with CRISPRa and the control gRNA, but treated only with the vehicle, served as an additional control **(Fig 6g)**. MEA analysis revealed that CRISPRa-mediated *NeuID* overexpression rescued the Aβ42-mediated defects in electrical activity **(Fig 6h)**, NAS **(Fig 6i)**, mean firing rate **(Fig 6j)** and frequency of network bursts **(Fig 6k)**.

Collectively, these findings suggest that increasing *NeuID* expression could serve as a novel therapeutic strategy for AD.

### NeuID interacts with EZH2 methyltransferase and regulates the expression of Olig2 transcription factor

Finally, we wanted to elucidate the mechanisms by which NeuID regulates neuronal function. Previous data have shown that nuclear lncRNAs can contribute to gene expression control by binding to regulatory elements on DNA, such as promoter regions, while simultaneously interacting with transcriptional regulators, including chromatin-modifying enzymes (*44*) (*45*) (*25*). Through this mode of action, lncRNAs can either recruit transcriptional regulators to specific genomic locations or act as molecular decoys (*46*) (*47*) (*25*) (*48*). To explore if such a mode of action may apply to *NeuID*, we employed Enrichr, an integrative tool that compiles multiple gene-set libraries to identify gene-regulatory mechanisms (*49*). This analysis revealed that genes deregulated upon *NeuID* knock down are likely controlled - at least in part - by EZH2. EZH2 is a subunit of the Polycomb Repressive Complex 2 (PRC2) that mediates gene-repression via histone 3 lysine 27 trimethylation (H3K27me3). Previous studies have shown that lncRNAs can recruit PRC2 to specific chromatin locations to silence gene-expression (*25*) (*50*). Moreover, reduced EZH2 function in the adult brain has been linked to the loss of neuronal identity and neurodegenerative phenotypes (*51*). EZH2 was shown to bind the lncRNA HOX antisense intergenic RNA (*Hotair)* (*52*). The identified binding region contains short guanine repeats that fold into guanine quadruplexes (G-quadruplexes) which are essential for RNA-protein interactions (*52*). A bioinformatic analysis revealed that the EZH2 binding site detected within *Hotair* is conserved in both the mouse and human variants of *NeuID* **(Fig. 7b)**, suggesting that *NeuID* may interact with the PRC2 complex and more specifically with EZH2. To test this, we performed RNA immunoprecipitation using an EZH2-targeting antibody. This experiment confirmed that *NeuID* binds to EZH2 **(Fig. 7c)**. In the case of *Hotair* it was shown that its binding to EZH2 regulates its ability to maintain H3K27me3 thereby helping to maintain gene-expression control (*53*). Therefore, we hypothesized that KD of *NeuID* might alter neuronal H3K27me3 levels, which could help explain the increased expression of non-neuronal genes observed upon *NeuID* KD. To explore this hypothesis, we performed H3K27me3 ChIP-sequencing on primary hippocampal neurons treated with either control or *NeuID* Gapmers. Principal component analysis (PCA) revealed that neuronal H3K27me3 patterns are affected by *NeuID* KD **(Fig. 7d)**.

**Figure 7.**
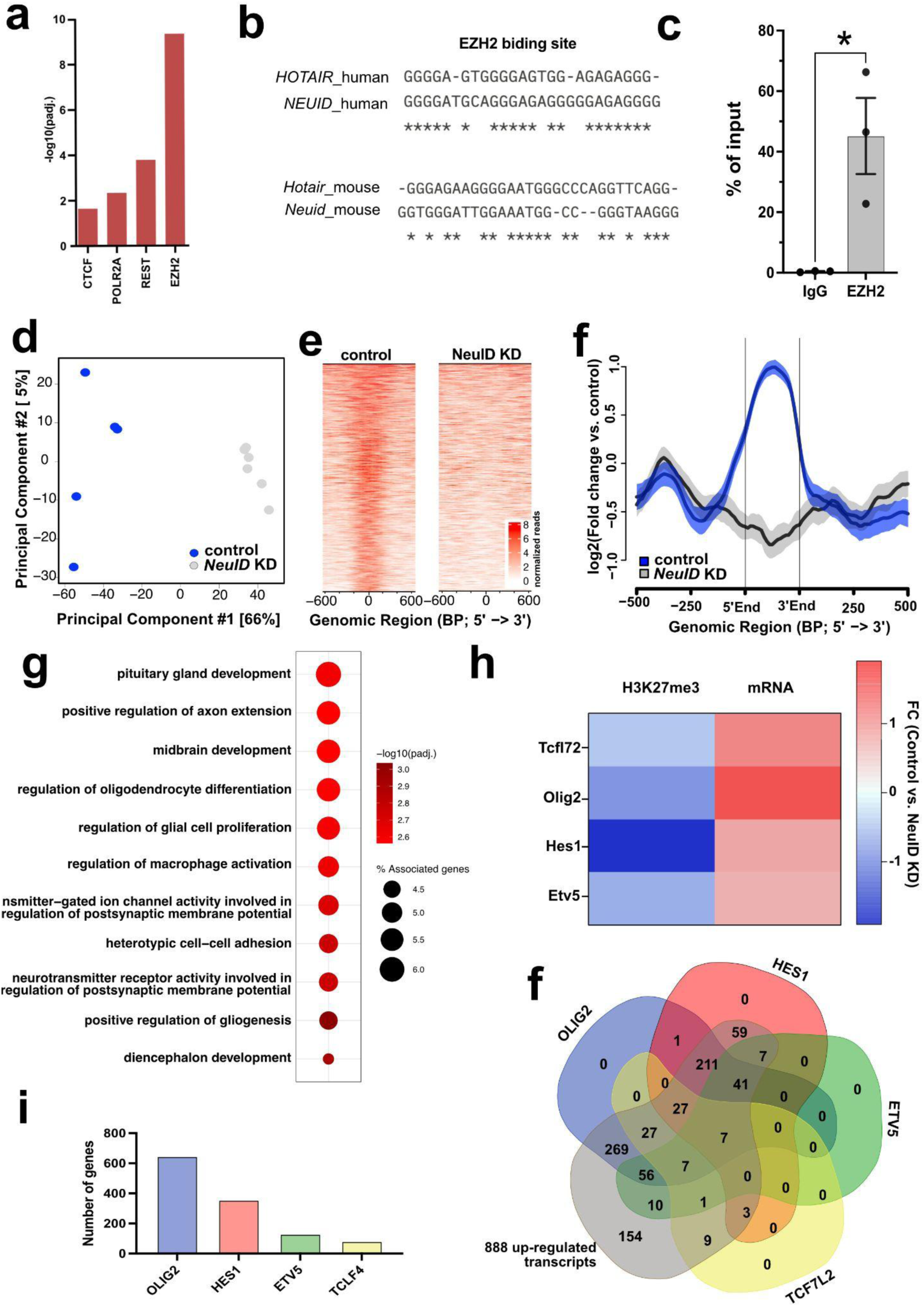
*NeuID* interacts with the EZH2 and affects H3K27 methylation of non-neuronal genes in neurons. **a.** EnrichR analysis using the genes de-regulated upon *NeuID* knock down as input reveals *EZH2* as a potential interaction partner for *NeuID*. **b.** Sequence comparison of the reported *EZH2* binding region within the lncRNA *Hotair* to *NeuID* in mouse and humans reveals strong similarity. **c.** Bar graph depicting the results from RNA-immunoprecipitation showing the interaction of *NeuID* with *EZH2*. IgG was used as control. unpaired tTest *p<0.05. **d.** Principal component analysis showing the results for H3K27me3 Chip-seq in neurons upon *NeuID* knock down in comparison to control (n =5/group) **e.** Heat map and **(f)** NGS plot showing the distribution of H3K27me3 along genes down-regulated upon *NeuID* knock down showing significantly altered H3K27me3 levels. **g.** Dot blot showing significantly enriched GO terms from the genes depicted in **(e)** and **(f)**. **h.** Heat map showing H3K27me Chip-seq and mRNA levels detected via RNA-seq signals for key transcription factors normally specific to glial cells. Depicted is the log2FC value. **i.** Bar chart showing the number of genes upregulated upon NeuID knockdown that are predicted targets of each of the displayed transcription factors. **f.** Venn diagram showing the overlap of genes upregulated upon NeuID knockdown that are predicted targets of each of the displayed transcription factors. Note that out of 888 upregulated genes, only 154 could not be explained by the four transcription factors. Error bars show SEM.

Next, we performed a differential binding analysis to identify H3K27me3 peaks that significantly differed between control and *NeuID* KD neurons (**see Supplementary Table 5**), and intersected these with transcripts that were significantly upregulated upon NeuID knockdown.

Through this approach, we identified 62 genes that had completely lost H3K27me3 occupancy and were significantly upregulated following *NeuID* KD **(Fig. 7 e, f)**. GO term analysis of these genes revealed that they represent developmental (“pituitary gland development”, “midbrain development”,” diencephalon development”) and glial cell related processes (“regulation of oligodendrocyte differentiation”, “regulation of glial cell proliferation”, “positive regulation of gliogenesis”) **(Fig. 7g, supplemental table 6)**. Upon closer inspection of the genes that exhibit decreased H3K27me3 binding and increased expression upon *NeuID* knock down, we detected 4 key transcription factors that are specific to glial cells under physiological conditions, namely *Etv5, Hes1*, *Olig2*, and *Tcfl2* **(Fig. 7h).**

While the 62 genes exhibiting reduced H3K27me3 occupancy and increased expression upon *NeuID* knockdown are likely directly regulated via *NeuID* acting through the PRC2 complex, we hypothesized that the increased expression of the four transcription factors may help explain — at least in part — the differential expression of the remaining genes that were upregulated following *NeuID* knockdown. To test this, we performed a bioinformatic analysis to determine how many of the 888 genes upregulated upon *NeuID* knockdown contained promoter binding sites for ETV5, HES1, OLIG2, and/or TCF7L2. The data revealed that the majority of these genes could be regulated by OLIG2 (646), followed by HES1 (356), ETV5 (129), and TCF7L2 (82) **(Fig. 7i)**. A combined analysis of the predicted target genes of the four transcription factors showed that 734 (84%) of the 888 upregulated genes could potentially be explained by increased activity of these factors **(Fig 7f, supplemental table 7)**.

## Discussion

In this study, we report the discovery and first functional characterization of *NeuID*, a previously unstudied, neuron-specific, and brain-enriched lncRNA. Through integrative transcriptomic analysis across human and mouse datasets, we identified *NEUID* as one of the few lncRNAs that is conserved, enriched in the brain, and selectively expressed in neuronal populations. We have previously used this approach to identify lncRNAs in other cell types relevant to AD pathology. For example, we characterized the astrocyte-specific lncRNA *PRDM16-DT* and demonstrated its functional relevance in AD (*25*). These findings are in line with previous data reporting roles for lncRNAs in AD pathogenesis such as *BACE1-AS*, *BDNF-AS* or *MEG3* (*14*) (*54*) (*55*) (*11*) (*8*) are likely to support further discovery of lncRNAs involved in neurodegeneration, emphasizing the utility of our strategy in exploring the lncRNAome for therapeutic targets.

*NEUID* is significantly downregulated in three independent cohorts of AD brain samples, and its expression inversely correlates with B&B staging, suggesting a role in early disease progression. Importantly, *NeuID* levels are also reduced in response to Aβ42 *in vitro*, under conditions that do not readily induce neurodegeneration, while *NEUID* levels are not altered in FTD, indicating that *NEUID* loss is not merely a secondary effect of neuronal cell death.

Mechanistically, our data suggest that *NeuID* operates through an epigenetic axis involving the PRC2 complex. We show that *NeuID* interacts with EZH2, affecting H3K27me3 levels at loci associated with developmental and glial-specific genes. This is consistent with previous studies demonstrating that the lncRNA *Hotair* recruits EZH2 to specific chromatin regions to direct H3K27me3 deposition (*52*) (*53*). Supporting a similar mechanism for human and mouse *NeuID*, we identified a conserved EZH2-binding motif within its sequence, homologous to the EZH2 interaction domain in *Hotair*.

Although additional binding partners cannot be excluded, our findings support the emerging hypothesis that cell-type–specific lncRNAs orchestrate the activity of ubiquitously expressed chromatin regulators like EZH2, thus maintaining cellular identity. This concept, previously proposed in other systems (*56*) (*57*) (*58*), is here extended to the context of neuronal identity and neurodegeneration. In *NeuID*-deficient neurons, we observed derepression of 888 genes, many involved in developmental or glial pathways, consistent with a loss of neuronal identity. Among them, 62 genes showed decreased H3K27me3 occupancy, suggesting direct regulation by *NeuID*. These 62 genes were enriched for developmental and glial-specific pathways, and notably included key transcription factors such as *Olig2*, *Etv5*, *Tcf7l2*, and *Hes1*, which are important for brain development and are linked glia cell function in the adult brain (*59*) (*60*) (*61*) (*62*). Our motif analysis indicates that these transcription factors may explain up to 85% of secondary transcriptional changes observed upon *NeuID* KD, further supporting the view that a key function of *NeuID* is to orchestrate the silencing of non-neuronal genes in neurons by guiding the PRC2 complex.

Furthermore, these data imply that *NeuID* functions as a molecular recruiter of PRC2 to the promoter of non-neuronal target genes. This contrasts with the recently described lncRNA *PRDM16-DT*, which interacts with PRC2 as a decoy, blocking its activity to preserve astrocyte identity. Together, these findings suggest that in the brain, one function of nuclear, cell-type–specific lncRNAs may be to either guide or prevent PRC2 activity to maintain the correct cell-type specific epigenetic landscape. Clearly more research is needed to further test this hypothesis for PRC2 and other chromatin modifying complexes.

Loss of *NeuID* not only altered gene expression but also disrupted neuronal physiology. We observed reduced dendritic spine density, fewer synapses, impaired neuronal network synchronization, and deficits in hippocampus-dependent memory, which was accompanied by decreased expressed of genes linked to neuronal plasticity. This is in line with previous studies showing that manipulation of lncRNAs can affect neuronal plasticity and memory function (*63*) (*64*) (*7*). Since we did not observe evidence that *NeuID* directly regulates these genes, we suggest that the loss of neuronal function and the downregulation of genes related to neuronal plasticity is a secondary effect—reflecting the consequences of neurons entering a developmental and glial cell–specific gene expression program.

Notably, *NeuID* was downregulated by Aβ42 exposure, a hallmark of early AD pathology, and CRISPRa-mediated overexpression of *NeuID* restored physiological expression levels and reinstated Aβ42-induced defects in neuronal network activity. To the best of our knowledge, this is the first time such an effect is described for a neuronal lncRNA, although previous studies have reported that lncRNAs can exhibit altered expression in models for amyloid pathology (*14*) (*65*) (*66*).

Although we show that *NeuID* regulates H3K27me3 via EZH2, the broader epigenetic impact of *NeuID* remains incompletely mapped. Many of the transcriptional changes observed upon *NeuID* KD are likely indirect. This view is supported by our finding that, out of the 888 upregulated genes, only 62 may be directly regulated by *NeuID* through its interaction with EZH2. Notably, among these 62 genes were key transcription factors that could account for up to 85% of the upregulated genes following *NeuID* KD. Most prominent was OLIG2, which is consistent with previous studies showing that elevated levels of OLIG2 in neural progenitor cells are linked to the pathogenesis of Down syndrome, an intellectual disability disorder associated with beta amyloid pathology (Chakrabarti, 2010} (*67*).

As this is the first report of *NeuID* there are several other key questions to be addressed in future research. For example, the mechanisms governing NeuID’s neuron-specific expression remain not well understood. It will be important to address this question not only for *NeuID*, but all brain and cell-type specific lncRNAs. In addition it will be important to elucidate how AD risk factors such as Aβ42 downregulate *NeuID* and the therapeutic potential for targeting *NeuID* needs to be tested in animal models *in vivo*. This is however, getting increasingly complicated, due to restricted animal care policies, at least within Europe, and Germany in particular. Thus, further research in additional AD model systems such as human iPSC derived organoids would be an important approach for future research.

From a translational perspective, it would be important to assess *NEUID* expression in liquid biopsies from patients at various stages of AD pathogenesis. The analysis of non-coding RNAs (ncRNAs) as biomarkers for Alzheimer’s disease is an active area of research, particularly in the case of microRNAs (*68*) (*69*) (*70*). However, emerging data also support the view that analyzing lncRNAs in blood may represent a promising strategy for the development of novel biomarkers in AD (*71*) (*72*). This could support the development of stratified therapies towards lncRNA expression and/or functionthat might be applied in combination with other approaches, such as anti-amyloid-based treatments.

In summary, our findings identify *NeuID* as a critical regulator of neuronal identity and plasticity, with direct implications for AD. Its strict neuronal specificity and defined mechanism of action position it at the intersection of epigenetic control, synaptic integrity, and memory function—making it a compelling and precise target for RNA-based therapeutic strategies in neurodegeneration.

## Methods

### Human tissues

For the analysis of *NEUID* expression by qPCR, brains (prefrontal cortex, BA9) from control (n = 2 females & 2 males; age = 81.75 ± 7.41 years, PMD = 13.75 ± 4.99 h) and AD patients (n = 2 females & 4 males; age = 85.83 ± 11.63 years, PMD = 8.5 ± 7.45 h; Braak & Braak stage IV) were obtained with ethical approval from the ethics committee of the University Medical Center Göttingen and upon informed consent from the Massachusetts Alzheimer Disease Research Center (NIA P50 AG05134, Boston, USA). *NEUID* expression data in human dorsolateral prefrontal cortex autopsy samples of the FHS participants are derived from previous publications (*30, 31*).

The same human left ventricular heart tissue (RNA was already extracted) as previously described (*73*) were used. The samples represent healthy human hearts unassigned for transplantation. All performed experiments conform to the Declaration of Helsinki and all patients provided written informed consent for the use of cardiac tissue samples.

### Animals

The protocol to generate primary neurons and to collect tissue from adult mice was approved by the Lower Saxony State Office for Consumer Protection and Food Safety. C57B/6J or CD1 mice (Janvier Labs, Le Genest St Isle, France) were kept under a 12h/12h light/dark cycle, in standard single cages with food and water provided ad libitum. For obtaining brain subregions, liver and heart (left ventricle) samples for nuclei extraction or direct RNA extraction, adult animals were sacrificed via cervical dislocation and the organs were immediately removed on ice, followed by liver, left ventricle and anterior cingulate cortex (ACC) dissection. Other selected brain subregions (cortex, as well as dentate gyrus, CA1 and CA3 regions of hippocampus) were dissected in ice-cold PBS under a dissection microscope. Acquired tissue was snap-frozen in liquid nitrogen and stored at -80°C until further processing. Behavioral experiment s and stereotactic injection were approached by the animal care committee of the Albert Einstein College of Medicine, Bronx, NY, USA.

### Fear conditioning

Contextual fear conditioning was performed in an automated system (TSE Systems) as previously described (*74*). Briefly, mice were exposed for 3 min to a novel context (A), followed by footshock (2 s, 0.7 mA, constant current) and tested 24 h later by re-exposing them to the same context. For trace fear conditioning, mice were exposed for 3 min to Context A followed by 3 consecutive pairings of tone (30 s, 75dB, 10kHz), temporal trace (15 s), and footshock (2 s, 0.7 mA, constant current). Memory tests were performed the next day by re-exposing the mice to Context B (3 min) followed by three presentations of tone (30 sec) and trace (15 sec) separated by 60-s inter-trial intervals (ITI) . Freezing, measured every 10 s, served as an index of memory. Freezing was expressed as a percentage of the total number of observations during which the mice were motionless. Activity was recorded automatically by an infrared beam system and expressed as cm/s. The individual experiments were not performed on littermates, so we did not apply randomization procedures, but all behavioral tests were performed by experimentalists who were unaware of the treatments because all injection solutions were coded by the lab technician.

### Nuclei isolation

The isolation of nuclei from mouse human brain tissue was performed according to our previously publications (*75*) (*28*) with minor modifications. Briefly, the tissue was homogenized using a dounce homogenizer in 500 µl EZ prep lysis buffer (Sigma NUC101) with 30 strokes (mouse tissue) or 60 strokes (Human tissue). The homogenate was transferred to 2 ml Eppendorf tube and the volume was adjusted to 2 ml with lysis buffer followed by 7 min incubation on ice. The homogenate was centrifuged for 5 min at 500 g at 4°C and the supernatant was discarded. The nuclei pellet was resuspended in 2 ml lysis buffer and incubated on ice for 5 min. The homogenate was centrifuged for 5 min (500 g at 4°C) and the pellet was resuspended in 1.5 ml nuclei storage buffer (NSB: 1 x PBS, 0.5% BSA, 1:200 RNAse inhibitor, 1:100 Roche protease inhibitor) followed by another centrifugation for 5 min (500 g at 4°C). The obtained pellet was resuspended in 1 ml NSB and stained with anti-NeuN-AlexaFlour® (catalog) for 1h at 4°C followed by centrifugation for 5 min (500 g at 4°C). The pellet was washed once with 500 µl NSB and resuspended in 300-500 µl NSB depending on the size of the pellet and stained with 1:100 7AAD (catalog).

The samples were passed through a 40 µm filter into FACS tubes and vortexed before sorting. Sorting of NeuN-positive and NeuN-negative nuclei was performed with FACS Aria III (BD Biosciences). Sorted nuclei were counted in the Countess II FL Automated counter. The sorted nuclei were then used for Single-nuclei RNA-seq or centrifuged and resuspended in TRIzol-ls for RNA isolation.

### Single nuclei total RNA sequencing and analysis

Single-nuclei RNA-seq was performed at the NGS Integrative Genomics Core Unit in Göttingen, Germany. Approximately 1200 single cells were sequenced per sample. Single nuclei total RNA-seq was performed as described in (*25*). In short, nuclei were first stained with Hoechst 33342 that enables selection of suitable nuclei for dispensing into the Takara ICELL8 5184 nano-well chip. CellSelect Software (Takara Bio) was used to visualize and select wells containing single nuclei. Four 5184 non-well chips were used for samples containing 1200 to 1400 nuclei/sample. Libraries were amplified and pooled as they were extracted from each of the single nanowell chip. Purification and size selection of libraries was done with Agencourt AMPUre XP magnetic beads (Beckman Coulter) to obtain average library size of 500 bp. Libraries were sequenced on the HiSeq 4000 (Illumina) to obtain 0.3 to 0.4 Mio reads per nuclei (SE; 50 bp). Bcl2fastq (v2.20.2) was used for converting BCL files to FASTQ format from raw images. Further processing was performed with the Cogent NGS Analysis pipeline (v1.5.1) to generate gene-count matrix. The demuxer (cogent demux) was used to create demultiplexed FASTQ files from the barcode sequencing data. The resulting output was then used as input for the analyzer (cogent analyze) which performs trimming, mapping, and gene counting to create a gene count matrix. Quality control was done by evaluating the quality report provided by the Cogent analyzer.

The SCANPY package was used for pre-filtering, normalization and clustering. First, cells with low quality were excluded. Then counts were scaled by total library size and transformed to log space. Highly variable genes were identified based on dispersion and mean, the technical influence of the total number of counts was regressed out, and the values were rescaled. Principal component analysis (PCA) was performed on the variable genes, and UMAP was run on the top 50 principal components (PCs). The top 50 PCs were used to build a k-nearest-neighbours cell-cell graph with k=50 neighbours. Subsequently, spectral decomposition over the graph was performed with 50 components, and the Leiden graph-clustering algorithm was applied to identify cell clusters. We confirmed that the number of PCs captures almost all the variance of the data. For each cluster, we assigned a cell-type label using manual evaluation of gene expression for sets of known marker genes by plotting marker gene expression on the UMAP and visual inspection. Violin plots for marker genes were created using the ‘stacked_violin function’ as implemented in SCANPY.

### Primary hippocampal neuronal culture

Primary hippocampal neuron cultures were prepared from CD1 mouse embryonic day 17 (E17) embryos. Briefly, a pregnant CD1 mouse was sacrificed using pentobarbital and the brains of the embryos were isolated, meninges were removed, and the hippocampi were dissected. To obtain single-cell suspension the tissue was digested with 2.5% trypsin followed by DNAse treatment. Cells were plated at a density of 60,000 cells/cm^2^ in poly D-lysine coated multi-well plates and grown in Maintenance media [Neurobasal with 1xB27 and 1mM GlutaMax]. Primary hippocampal neurons were used for experiments at DIV10-12.

### Antisense LNA Gapmers

Knockdown of NeuID in PHN was performed by using Antisense LNA gapmers packaged into Lipid nanoparticles (LNPs). The NeuID targeting and negative control (NC) gapmer was designed and purchased from Qiagen. The gapmers were packaged into LNPs using the Neuro9™ siRNA Spark™ Kit (5 nmol) with NanoAssemblr™ Spark™ system (Precision Nanosystems, Canada) according to the manufacturers instructions. Primary hippocampal neurons were transfected with NeuID targeting or NC LNPs on DIV 7 and the experiments were performed on DIV10.

### NeuID overexpression with CRISPRa

PHN were transfected with AAV-PHP.eB serotype for NeuID overexpression (OE). AAV vector containing the guide RNA (gRNA) and the dCas9-VP64 expression vector were purchased from Vectorbuilder.

PHN were co-infected with the gRNA containing AAV and the dCas9-VP64 expressing AAV at different multiplicity of infection (MOI) at DIV 7 to determine the optimum MOI. Experiments were performed 48h after virus infection.

### Abeta Treatment of Primary hippocampal neurons

Aβ (1–42) was purchased from AnaSpec. Aβ oligomers were prepared as previously described (*25*). Briefly, Aβ (1–42) was initially dissolved to 1mM in hexafluoroisopropanol (HFIP) and aliquoted. HFIP was removed by allowing evaporation under laminar hood. The peptide film was then stored desiccated at -20°C.

For aggregation, the peptide was resuspended in dry DMSO to a concentration of 5 mM. To obtain Aβ oligomers, Ham’s F-12 was added to bring the peptide to a final concentration of 100 μM and incubated at 4°C for 24 h. To assess the effect of Aβ oligomers on the expression of NeuID, primary hippocampal neurons were incubated with Aβ at mentioned concentrations for 24 h.

### RNAscope

RNAscope fluorescent multiplex assay (ACD Bio) combined with immunofluorescence for fresh frozen tissue sections was performed according to the manufacturer’s instructions and as described in (*25*). Briefly, sections were fixed in 10% neutral buffered formalin (NBF), dehydrated with ethanol, and treated with hydrogen peroxide followed by ON incubation at 4°C with anti-NeuN (catalog). The following day, in accordance with the RNAscope® Multiplex Flourescent Reagent Kit v2 (ACD Bio) protocol, probes designed against NeuID (ACD Bio) and TSA Plus Cyanine 5 (Akoya Biosciences, 1:1750) for detection were used. Afterwards, the sections were stained with Alexa Flour^TM^ 555 goat anti-rabbit secondary antibody (1:1000, ThermoFischer) and DAPI (Sigma-Aldrich). Images were acquired within a week after staining.

### Dendritic spine labelling

Dendritic spine density analysis on primary hippocampal neurons was performed as described previously (*76*). The cells were fixed with 2% paraformaldehyde (PFA) and washed 3x with PBS. For staining, 2-3 crystals of the DiI stain (Life technologies-Molecular Probes) were added to each well and incubated on a shaker for 10 min at RT. The cells were then washed until there were no visible DiI crystals in each well and incubated overnight at RT in PBS. Dendritic spine density was measured using dendritic spine counter plugin in ImageJ software.

### Synapse quantification

Synapse density quantification was performed by using a dual-color staining of pre- and post-synaptic markers along with staining of the neuronal body. The cells were fixed with PFA (4%, 10 min, RT) and permeabilized with 1% Triton X-100 in PBS followed by incubation in blocking buffer (2% BSA in PBS) for 1h at RT. The cells were then incubated with primary antibodies - Syn, Psd95, Map2 (Synaptic Systems) in blocking buffer overnight at 4°C. After washing with PBS, the cells were incubated with secondary antibodies in blocking buffer for 1h at RT. Synapse density was calculated using synquantvid plugin in ImageJ software.

### Multielectrode array recording and data analysis

To record spontaneous activity from primary hippocampal neurons, the Maestro Edge^TM^ multiwell microelectrode array (MEA) system (Axion Biosystems) was used. The Cytoview MEA 24, with 24 wells and 16 electrodes per well was used for the recordings. Primary hippocampal neuron culture was performed as described in a previous section, and the cell pellet was suspended in Maintenance Media with 1 µg/ml laminin to a concentration of 12.5 × 10^6^ neurons/ml. 8 µl of the cell suspension was added to each well of the MEA plate pre-coated with polyethylenimine. For recording, the MEA plate containing neurons was transferred to the recording chamber maintained at 37°C with 5% CO_2_, and the activity was recorded for 15 min. Data from the MEA was analyzed using the manufacturer’s analysis tool ‘Neural Metric Tool (Axion Biosystems)’.

### Imaging

All images were taken with the STEDycon STED/Confocal (Abberior) in the confocal mode at 63X oil immersion objective.

### RNA isolation, cDNA synthesis and RT-qPCR

Total RNA was isolated from cells or tissues using TRIzol (Invitrogen) and purified with RNA clean & concentrator-5 (Zymo research) according to the manufacturer’s instructions. RNA concentration was measured with nanodrop (ThermoFischer). Total RNA was reverse transcribed into cDNA using transcription first strand cDNA synthesis kit (Roche). RT-qPCR was performed on LightCycler® 480 system (Roche) using the SYBR Green I Master (Roche).

### RNA-seq and analysis

RNA quality was determined using Bioanalyzer (Agilent) prior to library preparation. Library preparation for RNA-seq was performed using SMARTer® Stranded Total RNA-seq kit v2-pico input according to the manufacturer’s instructions. Five ng of total RNA was used as input for library preparation. The quality and size of the library was determined using Bioanalyzer. The multiplexed libraries were sequenced in a HiSeq 2000 (Illumina) with a 50 bp single-read configuration.

The sequencing data was processed using a customized in-house software pipeline. Illumina’s conversion software bcl2fastq (v2.20.2) was employed for adapter trimming and converting the base calls in the per-cycle BCL files to the per-read FASTQ format from raw images. Quality control of raw sequencing data was carried out using FastQC (v0.11.5) (http://www.bioinformatics.babraham. ac.uk/projects/fastqc/). Reads were aligned using the STAR aligner (v2.5.2b) and read counts were generated using featureCounts (v1.5.1). The mouse genome version mm10 was utilized. Differential gene expression analysis was performed with DESeq2 (Version 1.42) (*77*) with RUVseq to correct for unwanted variation. GO term analysis was performed with ClueGO plugin in Cytoscape. Analysis for the enrichment of transcription factors was performed using ENRICHR (https://maayanlab.cloud/Enrichr/).

### Promoter region analysis

The promoter regions for 888 genes upregulated after loss of *NeuID* were obtained from the Ensembl database using Mus musculus genome GRCm39. Promoter regions were defined as 1000bp before TSS. The sequences were extracted using SAMtools (v1.10). Binding motifs for the glia-specific transcription factors were taken from the JASPAR (v2022) database. We used BLAMM (v1.0.0) to screen promoter regions for TF binding motifs.

### RNA immunoprecipitation

The nuclear and cytoplasmic fractions were obtained from cultured PHN using Nuclei isolation kit (Sigma) according to manufacturer’s instructions. The obtained nuclear fraction is resuspended in TSE buffer (10 mM Tris, 300 mM sucrose, 1 M EDTA, 0.1% Nonidet P-40, 100 units/ml RNAse inhibitor, 1x protease inhibitor), transferred to bioruptor tubes (Diagenode) and sonicated in a Bioruptor Plus (Diagenode) for 5 cycles (30s on, 30s off). The samples were then incubated on ice for 20 min with occasional vortexing and centrifuged for 10 min at 13000 rpm at 4°C. The supernatant was transferred to fresh tube and flash frozen at -80°C.

The lysate was precleared using 25 µl Pierce^TM^ Protein A/g magnetic beads (ThermoFischer) for 1h at 4°C to reduce unspecific binding. Anti-EZH2 (Millipore) or IgG isotype control (Millipore) was incubated with 50 µl Protein A/G magnetic beads for 2h at RT and washed with RNA-IP buffer (50 mM Tris-HCl, 100 mM NaCl, 32 mM NaF, 0.5% NP-40). 10 % of the lysate was kept as input and the precleared beads were added to the remaining lysate and incubate ON at 4°C on a rotator. Then the beads were washed 5x with RNA-IP buffer, and incubated in proteinase K for 1h at 37°C. The supernatant was collected and RNA extracted using the RNA clean & concentrator-5 kit (Zymo research).

### Chromatin immunoprecipitation

Primary hippocampal neurons were cultured in 6-well plates and treated with control or *NeuID* targeting LNPs as described above. To each well 1 ml of Low sucrose buffer (0.32 M sucrose, 5 mM CaCl_2_, 5 mM Mg(Ac)_2_, 0.1 mM EDTA, 10 mM HEPES, 0.1 % Triton X-100, 1 mM DTT, 1x Protease inhibitor) was added and the cells were collected from each well using a cell-scraper and immediately flash frozen in liquid nitrogen. To prepare chromatin for ChIP, the cells were thawed and cross-linked using 5% PFA for 10 min. The reaction was quenched by adding 125 mM glycine for 5 min followed by centrifugation at 1000 g for 3 min at 4°C. The supernatant was discarded and the nuclei pellet was resuspended in 500 µl Nelson buffer (140 mM NaCl, 20 mM EDTA, 50 mM Tris pH 8.0, 0.5% NP-40, 1 % Triton X-100, 1x protease inhibitor). The nuclei suspension was then homogenized using mechanical homogenizer (Ultraturax T10) with power on 4 for 10s. The suspension was then centrifuged at 10,000 g for 2 min at 4°C, and the pellet was washed once with Nelson buffer. Afterwards, the supernatant was discarded and the nuclei pellet was weighed and resuspended in 10x volume of RIPA buffer (140 mM NaCl, 1 mM EDTA, 1 % Triton X-100, 0.1% sodium deoxycholate, 10 Mm Tris-Cl, 1 % SDS, 1x Protease inhibitor). The samples were then incubated at 4°C on a wheel for 10 min and then sonicated for 30 cycles (30s on, 30s off). To check chromatin shearing a small amount of chromatin was aliquoted and crosslinked by RNAse A and proteinase K treatment for 1h at 65 °C. DNA was isolated using Zymo ChIP clean and concentrator kit (Zymo research) according to manufacturer’s instructions. The size of the sheared chromatin was determined by using Bioanalyzer 2100 with a DNA high sensitivity kit and the concentration was measured using Qubit 2.0 fluorometer (DNA high sensitivity kit). 1 µg of chromatin and 3 µg of antibody was used for h3k27me3 (Millipore) or IgG isotype control (Millipore).

### ChIP sequencing and analysis

The ChIP DNA was used for library preparation using NEBNext Ultra II DNA library preparation kit. Libraries were sequenced for single end 50 base pair reads with the NextSeq 2000 (Illumina). Base calling and fastq conversion of sequencing data was performed using the Illumina pipeline and quality control was performed using fastqc. Reads were mapped to mm10 mouse reference genome using bowtie2. MACS2 (*78*) was used for calling peaks with the option ‘broadpeaks’ and q value < 0.1. ngsplot (*79*) was used to create profile plots. Diffbind package was used for differential binding analysis with in-built DESEQ2 option for differential analysis (Ross-Innes, 2012). ChIPSeeker package of Bioconductor (Yu, 2015) was used for annotation of peaks.

### Stereotactic surgery and injection for behavior experiments

Double-guided cannulas were implanted in the dorsal hippocampus (DH) as described previously (*74*). Mice were anesthetized with 1.2% tribromoethanol (vol/vol, Avertin) and implanted with bilateral 26-gauge cannulas using a stereotaxic apparatus. Stereotaxic coordinates for the dorsal hippocampus were 1.8 mm posterior, ±1.0 mm lateral and 2.0 mm ventral to bregma, according to the mouse brain atlas (*80*). The *NeuID* Gapmer was stored as a 400 µM solution at -80oC, and, before use, heated at 65oC for 10 min, cooled on ice, and diluted 1:10. Each mouse received 20 pmol (10 pmol/0.25 µl/side) of NeuID or control (scramble oligonucleotide) .

### Statistical analysis

Statistical analysis was done using GraphPad Prism version 9. All graphs are represented as mean ± SEM unless stated otherwise. A two-tailed unpaired t-test or one-way ANOVA was used for analysis unless otherwise stated.

### Datasets used in this study

The following snucRNAseq datasets were used in this study. The human snucRNAseq datasets were from Schröder et al. (*25*). The scucRNAseq data from the mouse brain are from (*28*).

## Acknowledgments

AF was supported by the DFG (Deutsche Forschungsgemeinschaft) SFB1286 and GRK2824;The German Federal Ministry of Science and Education (BMBF) via the ERA-NET Neuron project EPINEURODEVO; The EU Joint Programme-Neurodegenerative Diseases (JPND) – EPI-3E; Germany’s Excellence Strategy - EXC 2067/1 390729940 and FS was supported by the GoBIO project miRassay (16LW0055) by the German Federal Ministry of Science and Education. AF, ID, JKB and TDS are supported by NIH RF1AG078299. TDS and JKB are supported by NIH grants P30AG072978 and U19AG068753. JKB is supported by NIH grant RF1AG078299 and TDS receives funding from the National Heart, Lung and Blood Institute (N01-HC-25195, 75N92019D00031 and HHSN268201500001) and the United States Department of Veterans Affairs, Veterans Health Administration, BLRD Merit Award I01BX005933 (TDS). RP, MSS and ED are students of the IMPRS Neuroscience programme (University Göttingen) and SS is a student of the IMPRS Genome Science Programme (University Göttingen).

## Author contribution

RP conducted most of the experiments, anylzed data and generated figures; ZP and JR performed stereotactic injection and performed behavioral experiments; MSS and IF contributed to the initial discovery of NeuID; SS performed RNAscope, DMK and TP contributed to the bioinformatic analysis; ED performed DIL dye staining, TDS, KB and ID analyzed the FHS data and ID provided postmortem brain tissue; KT provided human heart biopsies and SB, ALS and FS performed RNAseq experiments, AF conceived and supervised the project.

